# Multi-scale modeling reveals angiogenesis-induced drug resistance in brain tumor and predicts a synergistic drug combination targeting EGFR and VEGFR pathways

**DOI:** 10.1101/394668

**Authors:** Weishan Liang, Ji Zhang, Xiaoqiang Sun

**Affiliations:** Zhong-shan School of Medicine, Sun Yat-Sen University, Guangzhou 510080, China; Key Laboratory of Tropical Disease Control (Sun Yat-Sen University), Chinese Ministry of Education, Guangzhou 510080, China.; School of Mathematics, Sun Yat-Sen University, Guangzhou 510275, China.; Department of Neurosurgery, State Key Laboratory of Oncology in South China, Sun Yat-Sen University Cancer Center, Collaborative Innovation Center for Cancer Medicine, Guangzhou 510275, China.

**Keywords:** Multi-scale modeling, Angiogenesis, EGFR signaling pathway, VEGFR inhibition, Drug combination

## Abstract

Experimental studies have demonstrated that both the extracellular vasculature, microenvironment and intracellualr molecular network (e.g. epidermal growth factor receptor (EGFR) signaling pathways) are essentially important for brain tumor growth. Some drugs have been developed to inhibit the EGFR signaling pathways. However, how does angiogenesis affect the response of tumor cells to the drug treatment has rarely been mechanistically studied. Therefore, a multiscale model is required to investigate such complex biological systems that contain interactions and feedbacks among multi-levels. In this study, we developed a single cell-based multi-scale spatio-temporal model to simulate more realistic vascular tumor growth and drug response, based on VEGFR signaling pathways, EGFR signaling pathway and cell cycle as well as several microenvironmental factors that determine cell fate switches in a temporal and spatial context. The simulation reconstructed an evolving profile of vascular tumor growth, demonstrating the dynamic interplay between angiogenesis and various types of tumor cells (e.g., migrating, proliferating, apoptosis and quiescent cells). Moreover, we revealed the critical role of angiogenesis in the acquired drug resistance. We further investigated the optimal timing of combing VEGFR inhibition with EGFR inhibition and predicted that the drug combination targeting both EGFR pathway and VEGFR pathway has a synergistic effect. The experimental data validated the prediction of drug synergy, confirming the effectiveness of our model. The developed multiscale model explored mechanistic and functional mechanisms of angiogenesis underlying tumor growth and drug resistance, which advances our understanding of novel mechanisms of drug resistance and provides implications for designing more effective cancer therapies.

**Author summary:** Many targeted therapies have been designed to treat malignant tumors including gliomas, but the clinical effectiveness of these therapies are limited due to the emergence of drug resistance during cancer therapeutics. The mechanisms underlying cancer drug resistance have not been fully understood until now, which restricts the rational designing of robust and effective therapeutics. Increasing number of experimental studies have indicated that angiogenesis plays important role s in influencing the effect of drug treatment. However, how does angiogenesis affect the response of tumor cells to the drug treatment has rarely been mechanistically studied. In this study we developed a single cell-based multi-scale spatio-temporal model to investigate the role of angiogenesis in drug response of brain tumors. The model demonstrated dual roles of angiogenesis in drug treatment of brain tumors and revealed a novel mechanism of angiogenesis-induced drug resistance. Moreover, the model predicted a synergistic drug combination targeting both EGFR and VEGFR pathways with optimal combination timing. This study has been dedicated to elucidate mechanistic and functional mechanisms of angiogenesis underlying tumor growth and drug resistance, providing implications for designing more effective drug combination therapies.

## Introduction

Brain tumors, such as glioblastoma (GBM), are one of most malignant cancers with poor prognostic survival rate. Many targeted therapies have been designed to treat brain tumors, but the clinical effectiveness of these therapies are limited due to the emergence of drug resistance during cancer therapeutics. The mechanisms underlying cancer drug resistance have not been fully understood until now, which restricted the rational designing of robust and effective therapeutics. Therefore, it’s urgent to uncover the mechanisms of drug resistance for the success of more effective therapeutics of brain tumors.

A lot of experimental data have demonstrated that various factors are involved in the initiation and progression and drug response of brain tumors, ranging from genetic mutations, signaling pathways, to extracellular microenvironment and surrounding tissue. Previous studies of the drug resistance mechanisms have focused on the intracellular molecular scales, for instance, genetic/epigenetic mechanisms [1, 2], posttranslational modifications of proteins and reactivation of signaling pathways [3]. Recently, experimental studies have indicated that angiogenesis plays important role s in influencing the effect of drug treatment [4, 5]. However, how does angiogenesis affect the response of tumor cells to the drug treatment has rarely been mechanistically studied.

Brain tumor is a complex biological system that contain interactions and feedbacks among multi-levels, including molecular networks, cellular interactions, microenviromental factors and tissue vasculature. Moreover, the interactions among these scales are temporally evolving and spatially heterogeneous. Therefore, a multiscale dynamic spatio-temporal model is required to investigate the role of angiogenesis in tumor growth and drug response.

In the past decades, a variety of mathematical or computational models have been developed to simulate tumor growth and drug response. For example, ordinary differential equations (ODEs) models [6-8], stochastic process [9-12] or stochastic differential equations [13], partial differential equations [14], agent-based model [15-17] or cellular automata model [18, 19], and hybrid model [20, 21]. These models have been developed to describe cell population dynamics or to simulate microenvironment interactions and advanced our understanding of tumor progression and drug resistance. However, they fall short on integrating the effect of targeted drug therapies in particular of a vascularized tumor which include the interactions between tumor cells and angiogenesis as well as the related signaling pathways. In order to investigate the role of angiogenesis in the response of tumor cells to the EGFR-tyrosine kinase inhibitor (TKI) treatment that is used in clinical trials, it requires to integrate the drug treatment effects of EGFR inhibitor targeting tumor cells and VEGFR inhibitor targeting endothelial cells into a multiscale model of vascular tumor.

In this study, we extend our previous two-dimensional (2-D) multiscale agent-based model [22] to a more realistic three-dimensional (3-D) space and incorporate VEGFR inhibitor treatment based on its action mechanisms on VEGFR signaling pathways. Our model reconstructed an evolving profile of vascular tumor growth and demonstrated the dynamic interplay between various types of tumor cells (e.g., migrating, proliferating, apoptosis and quiescent cells) and the growth of blood vessels. With the incorporation of EGFRI treatment, the model revealed angiogenesis-induced drug resistance. Interestingly, it showed that the survival rate of tumor cells decreased in the early stage but rebound in a later stage. Moreover, inhibiting blood vessels’ growth using VEGFR inhibitor prevented the recovery of survival rate of tumor cells in the later stage, demonstrating the critical role of angiogenesis in the acquired drug resistance. We further investigated the optimal timing of combing VEGFR inhibition with EGFR inhibition and predicted that the drug combination targeting both EGFR pathway and VEGFR pathway has a synergistic effect. The experimental data validated the prediction, confirming the effectiveness of our model.

## Results

We first demonstrated the clinical relevance of angiogenesis-regulating VEGFR pathways and the related genes in glioma patients using clinical data. We then mechanistically modeled vascular tumor growth to understand the dynamic mechanisms of angiogenesis in cancer progression and drug response. We next investigated the role of angiogenesis in the drug resistance. Moreover, we examined the combination therapy using EGFR inhibitor and VEGFR inhibitor targeting both tumor cells and endothelial cells. Furthermore, we used experimental data to validate the model predictions of drug synergy.

### Clinical relevance of angiogenesis pathways in glioma patients

VEGFR signaling pathway regulates endothelial cell’s survival, proliferation and migration during angiogenesis through PI3K/AKT, PKC/ERK, and FAK/p38 pathways [23, 24] (Fig 1A). We examined whether VEGF and VEGFR genes, as well as other genes in VEGFR signaling pathways correlated to the survival of glioma patients. We analyzed clinical survival data and RNA-seq data of glioma patients that were downloaded from TCGA database. COX PH model was used to compute the risk score predicted by expression of VEGF and VEGFR genes (Fig 1B) or by expression of all collected genes in VEGFR signaling pathways (Fig 1C). K-M curves (Fig 1B-C) demonstrated that the survival rates of high and low risk patients were significantly different, assessed using the log-rank test. These results indicated the genes in the VEGFR signaling pathways significantly associated to the disease progression of GBM patients.

**Figure 1.**
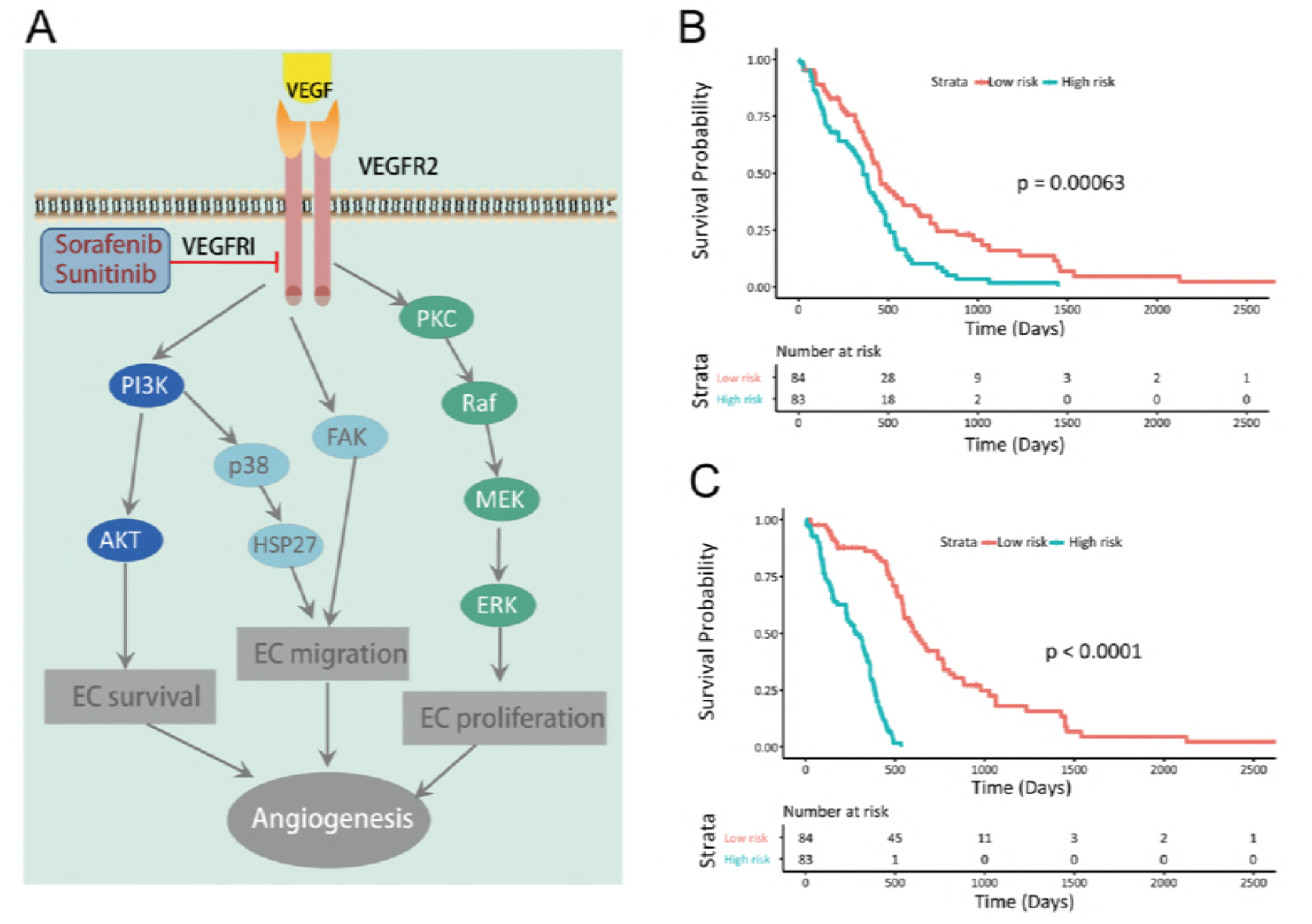
VEGFR signaling pathways in angiogenesis and the clinical associations of angiogenesis-related genes with survival rates of glioma patients. (**A**) The mechanism of VEGFR pathway regulating endothelial cells’ survival, proliferation and angiogenesis. TAFs such as VEGF can bind to its receptor, VEGFR and stimulate the signaling pathways including PI3K and ERK pathways, which regulates endothelial cells’ survival, proliferation and migration during angiogenesis [23, 24]. VEGFR inhibitor (VEGFRI), such as Sorafenib and Sunitinib, can inhibit the VEGFR signaling pathway by blocking VEGF-VEGFR binding. (**B**) Prognostic significance of VEGF and VEGFR genes. Shown are survival rates of glioma patients in low and high risk groups predicted by gene expressions of VEGF and VEGFR. (**C**) Prognostic significance of genes in VEGFR signaling pathways. Shown are survival rates of glioma patients in low and high risk groups predicted by expression levels of genes in VEGFR signaling pathways.

### Modeling vascular tumor growth and drug response

To understand the dynamic mechanisms of angiogenesis in cancer progression and drug response, we sought to mechanistically model vascular tumor growth in a more realistic situation. We developed a single cell-based multi-scale spatio-temporal model to simulate more realistic vascular tumor growth and drug response, based on VEGFR signaling pathways, EGFR signaling pathway and cell cycle as well as several microenvironmental factors that determine cell fate switches in a temporal and spatial context (Fig 2; See details in Methods section and **Text S1**). A novel algorithm was designed to simulate the VEGFR inhibitor effects on the blood vessels’ growth and integrated into the multiscale model of brain tumor, based on VEGFR signaling pathway and EGFR signaling pathway (Equations (1-6)).

**Figure 2.**
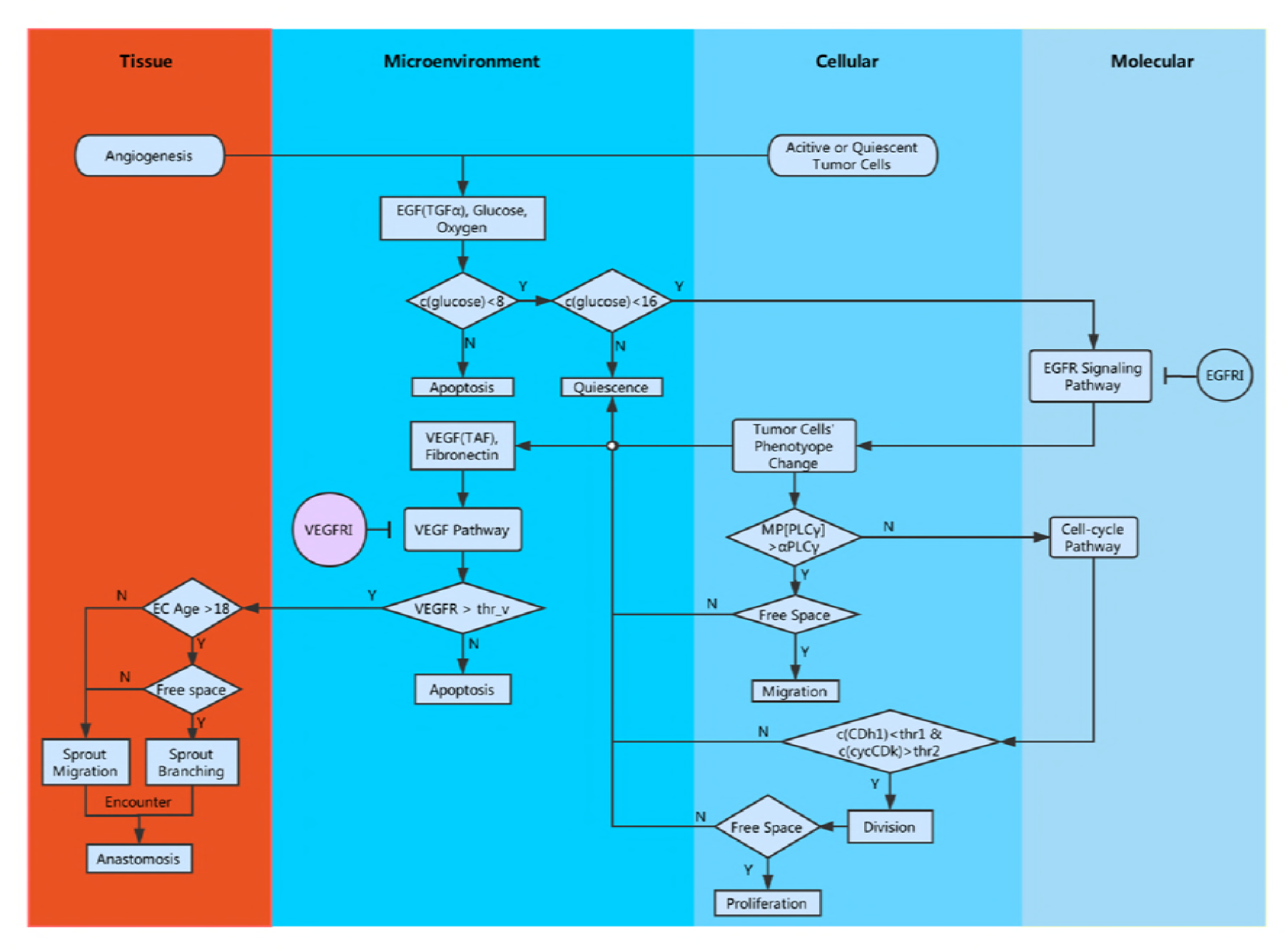
Flowchart of the computational modeling. Our model encapsulates four biological scales: molecular, cellular, microenvironmental and tissue scales. At molecular scale, EGFR signaling pathway, cell cycle and VEGFR signaling pathway were considered; at cellular scale, tumor cells switch their phenotypes and endothelial cells migrate, proliferate or dead; at microenvironment scale, growth factors, nutrients (glucose and oxygen) and chemical inhibitors diffuse and exchange; at tissue scale, new blood vessels grow and branch to form a micro-vasculature network. Intracellular signaling pathways were described using ODEs, and microenvironmental factors were described with PDEs. The cell’s phenotype switch was simulated using rule-based algorithm that is determined by both signaling pathways and microenvironmental factors. The treatment effects of EGFR inhibitor and VEGFR inhibitor were integrated into the model based on their action mechanisms of corresponding signaling pathways.

Fig 3 shows the vascular tumor growth profiles with and without EGFRI treatment in a 3-D space. Tumor cells were denoted in different colors according to their phenotypes: active (blue), proliferative (pink), quiescent (cyan) and apoptotic (black). The red lines at the bottom of the figures represent the blood vessels with several initialized tip endothelial cells. In the absence of EGFRI treatment (Fig 3A), the tumor cells grow more and more and generally develop into an olive shape with the increasing micro-vasculatures surrounded. While under the EGFRI treatment (Fig 3B), the tumor volume was much smaller than that without EGFRI treatment, showing an effective effect of EGFRI on repressing tumor growth at the early stage.

**Figure 3.**
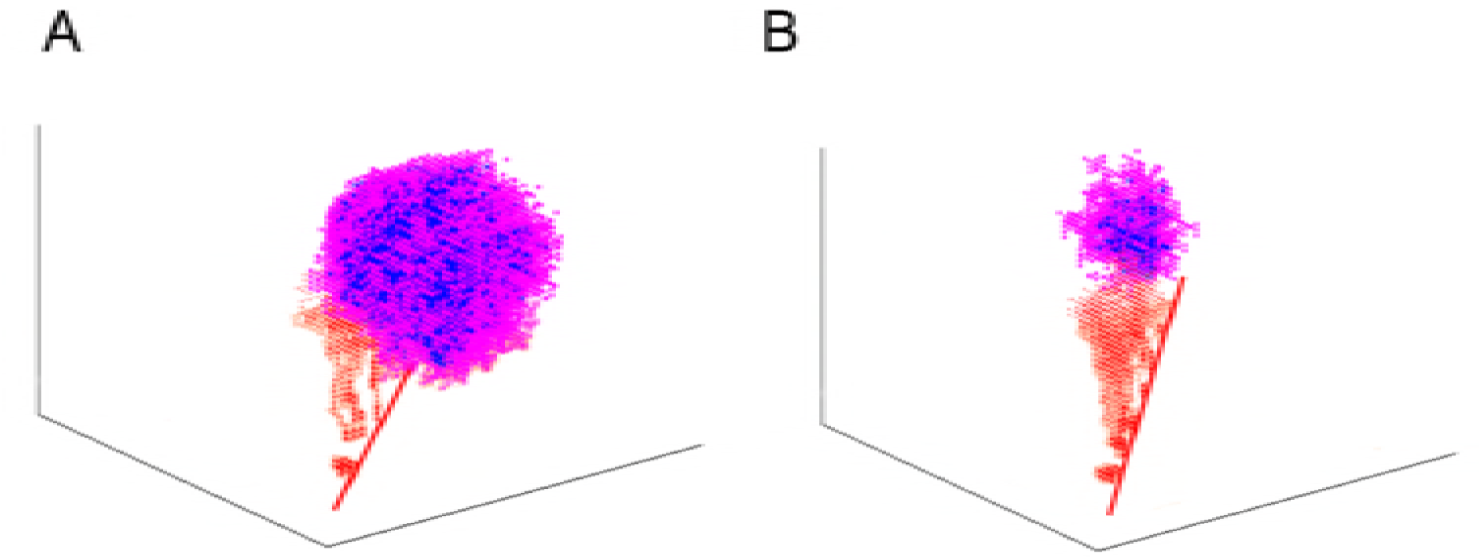
Vascular tumor pattern at 60 hours. (**A**) Vascular tumor growth pattern without EGFRI treatment. (**B**) Vascular tumor growth patterns under EGFRI treatment.

Fig 4(a) presents the evolution of number of various types of tumor cells and endothelial cells. The numbers of most cells changed with varying rates, showing nonlinearity of the tumor growth and heterogeneous evolutionary dynamics of various tumor cells. While the endothelial cells increased stably along time. Monitoring the whole process of vascular tumor growth, we found that after 110 hours, some apoptotic tumor cells appeared, and the proportion of apoptosis had getting increasingly larger as time went on until about 260 hours. After that, in response to the promoting effect of vasculatures, more and more tumor cells become active and move towards the denser micro-vasculatures. It is also noteworthy that the number of active tumor cells declined at around 70 hours and then increased after 250 hours. Besides, the proliferative cells’ amount fluctuated slightly at a relative low level. These results showed the dynamic phenotype switches between various types of tumor cells.

**Figure 4.**
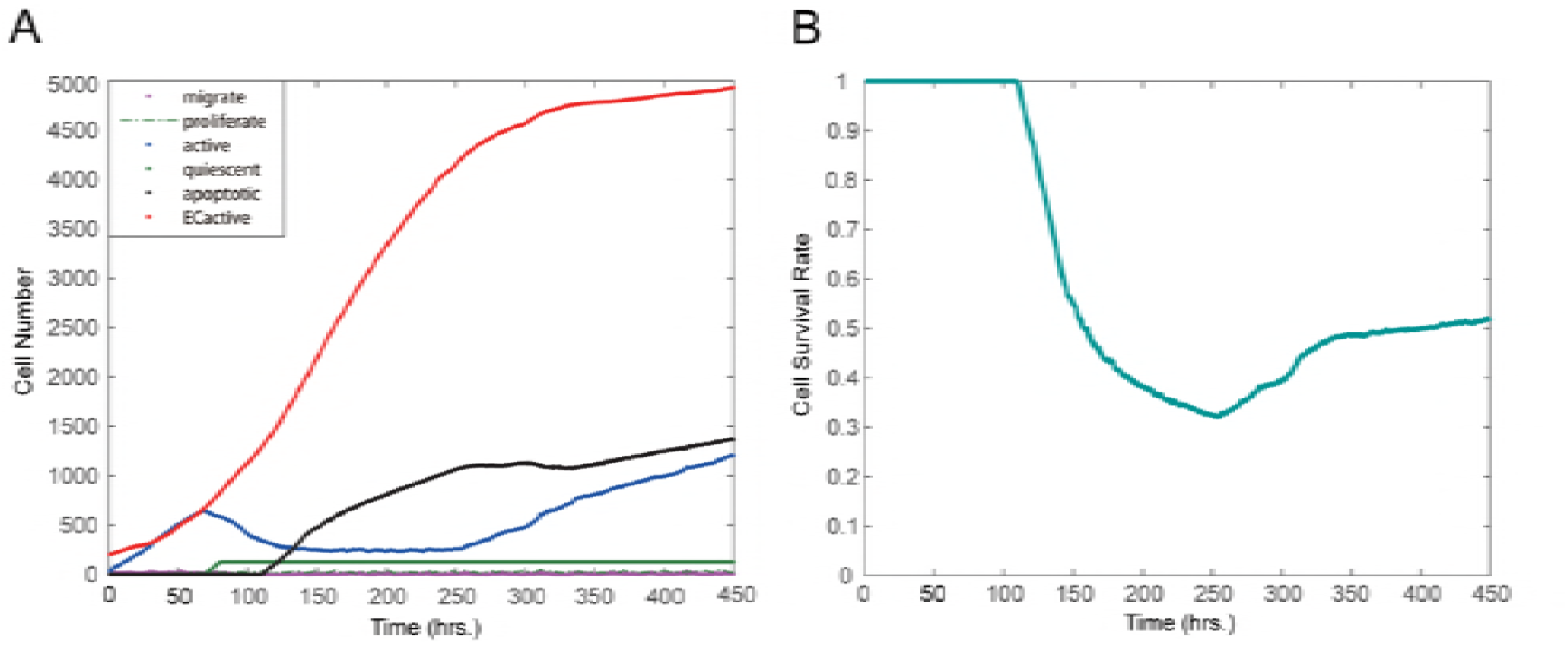
Changes of cell numbers and tumor survival rate under EGFRI treatment. (**A**) Changes in number of various cells under EGFRI treatment. (**B**) Changes in tumor cell survival rate.

We calculated the survival rate of tumor cells as a function of time. Fig 4(b) shows that, under EGFRI treatment, the survival rate of tumor cells showed a sharp drop at about 120 hours, and then kept decreasing rather slowly during 200 to 260 hours. However, at 260 hours, the rate rebounded, and reaches at approximately 52% at the end of the simulation (450 hours). This result demonstrated an emergence of drug resistance. As a consequence, the effect of EGFRI-only treatment was limited.

### Deciphering mechanisms of angiogenesis-induced drug resistance

Next we sought to identify the driving force of the drug resistance to EGFRI treatment. Considering the interplay between angiogenesis and tumor cells during EGFRI treatment as described above, we hypothesized that the angiogenesis played essential roles in tumor growth and drug response, which induced glioma resistance to EGFRI treatment.

To test the above hypothesis *in silico*, we integrated VEGFR inhibitor (VEGFRI) treatment effect into the model to suppress angiogenesis. Fig 5(a-d) shows vascular tumor growth patterns at different time points under the treatment of EGFR inhibitor combining VEGFR inhibitor at 240 hours. By adding VEGFRI treatment, angiogenesis was suppressed and the tumor growth was controlled after 240 hours, in contrast with the EGFRI-only treatment before 240 hours. Fig 5(e) shows changes in number of various cells in response to the combined VEGFRI and EGFRI treatment with VEGFRI added at 240 hours. The angiogenesis was inhibited after adding VEGFRI treatment. Meanwhile, the apoptotic tumor cells increased sustained and the active tumor cells kept in a low level.

**Figure 5.**
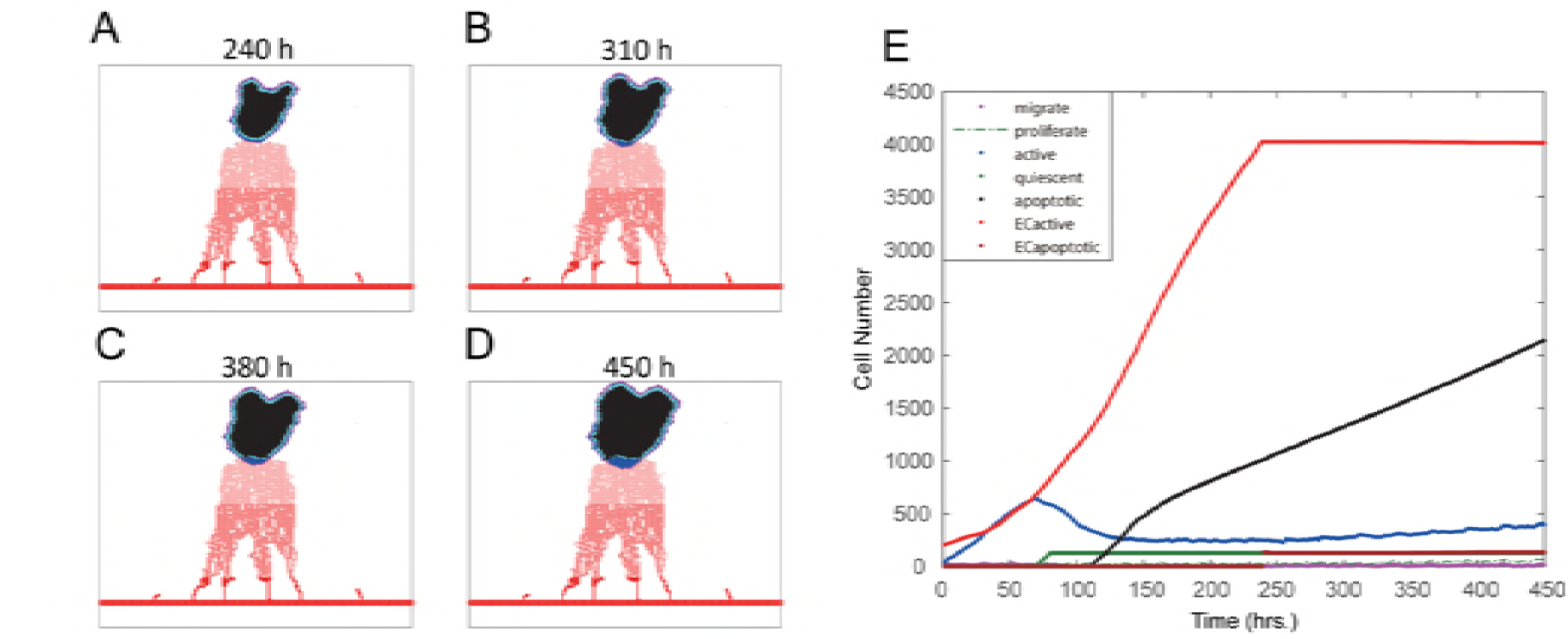
Vascular tumor growth patterns at different time points under the treatment of EGFR inhibitor combining VEGFR inhibitor at 240 hours. (**A**) Vascular tumor growth patterns at hour 240; (**B**) Vascular tumor growth patterns at hour 310; (**C**) Vascular tumor growth patterns at hour 380; (D) Vascular tumor growth patterns at hour 450. (E) Changes in number of various cells in response to the combined VEGFRI and EGFRI treatment. VEGFRI was added at 240 hours.

Fig 6a shows that adding VEGFRI at 240 hours (i.e. the turning point before survival rate curve) resulted in the sustained decrease of survival rate, in contrast to the survival rate without VEGFRI. Blocking angiogenesis indeed prevented the recurrence of tumor growth, suggesting the critical role of angiogenesis in driving drug resistance.

**Figure 6.**
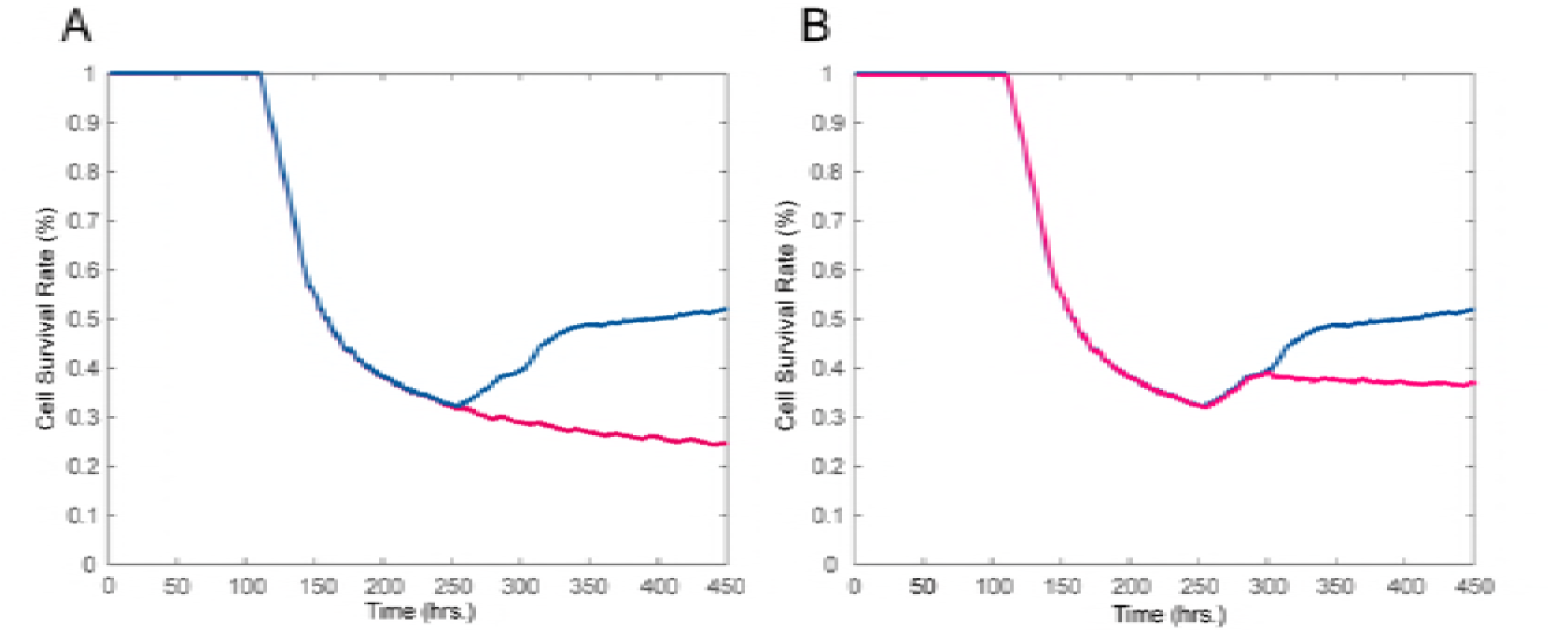
Effect of timing of combining VEGFR inhibition. (A) Combining VEGFRI treatment at hour 240; (B) Combining VEGFRI treatment at 300 hours. The blue and black lines represent the tumor cell survival rates with and without VEGFRI treatment, respectively.

These results revealed the dual roles of angiogenesis in the emergence and development of drug resistance of brain tumors to EGFRI treatment. More precisely, angiogenesis could delivery drugs or chemical inhibitors to the region of tumor cells (**Fig S3**), resulting in the increase of apoptosis and quiescent phenotype of tumor cells. While, on the other hand, the neo-vasculatures could transport nutrients such as glucose and oxygen (**Fig S4**) to tumor cells to maintain their survival and revisable phenotype switching to active or migrating status. Provided these dynamic interactions, the tumor cells’ survival rate rebounded at the later stage and drug resistance emerged.

### Synergistic combination of EGFRI and VEGFRI

Furthermore, we investigated the effect of different timing of combining VEGFRI treatment with EGFRI treatment on the emergence of drug resistance. We added VEGFRI at different time points, either before the turning point of survival rate curve (i.e. 260 hours) or after the emergence of drug resistance. We found that adding VEGFRI after 240 hours could also prevent the rebound of survival rate but resulted in a higher steady state compared to adding VEGFRI at 240 hours, as shown in Fig 6b. In addition, **Fig S1** demonstrated that adding VEGFI before 240 hours has almost the same effect on the survival rate with that of adding VEGFI at 240 hours. These results suggest that combining EGFRI with VEGFRI treatment could effectively rebound of tumor cells’ survival rate. Moreover, the optimal timing of adding VEGFRI treatment should be around 240 hours, slight earlier than the time point (260 hours) at which survival rate curve rebounded. Although adding VEGFRI more earlier before 240 hours had almost the same effect on preventing the rebound of the survival rate, adding drugs too early would produce more side effect.

Moreover, we used Bliss combination index to evaluate the combination effect between EGFRI and VEGFRI (See methods). We predicted that the drug combination targeting both EGFR pathway and VEGFR pathway has a synergistic effect, since the combination index is greater than 0 (Fig 7a). We used *in vivo* experimental data [25] of mice with brain tumors to validate this prediction. The calculation of the combination index for the experiments also resulted in synergistic effect of the combination of an EGFR inhibitor (DC101, 4mg/kg) and a VEGFR inhibitor (Cetuximab, 1mg/kg), as shown in Fig 7b. Therefore, the experimental data supported the model prediction, confirming the effectiveness of our model.

**Figure 7.**
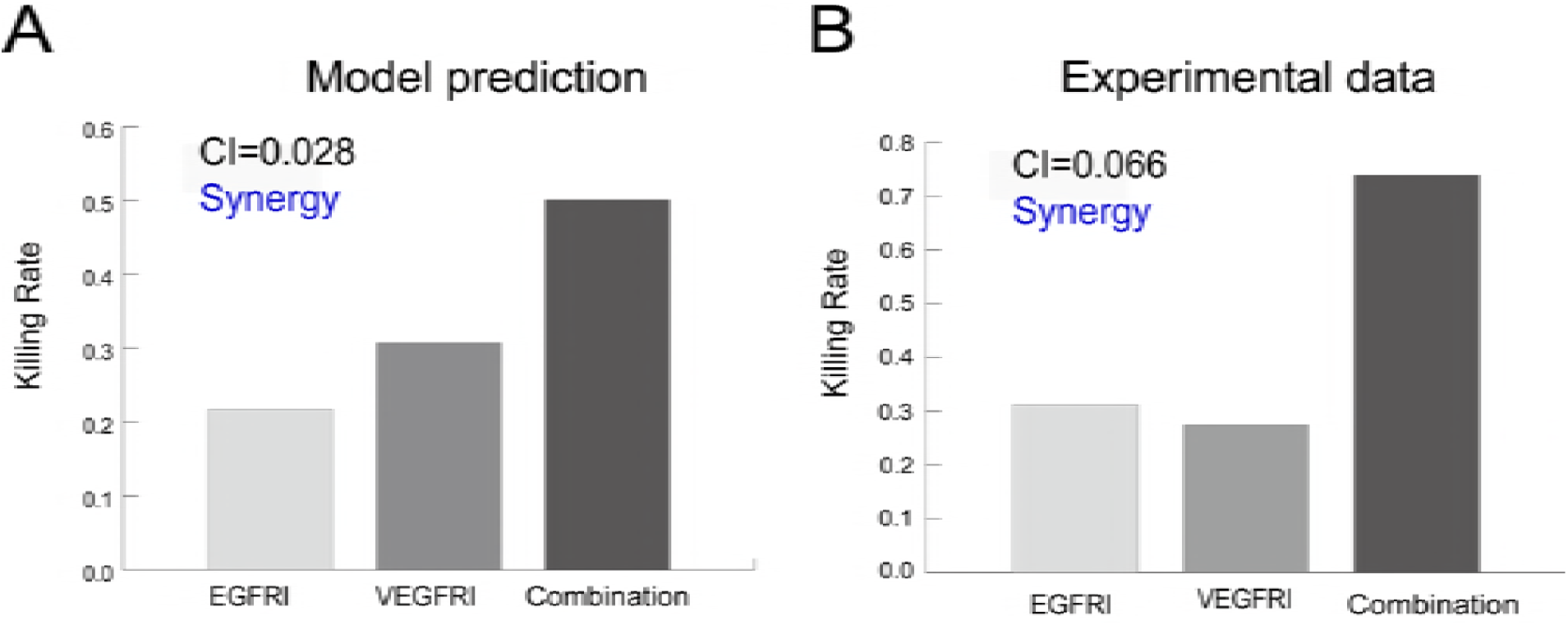
Validation of the synergistic effect of the drug combination targeting both EGFR and VEGFR. (**A**) Prediction of the synergistic effect of the drug combination targeting both EGFR and VEGFR. (**B**) Experimental validation using *in vivo* data of the DC101 (EGFR inhibitor), Cetuximab (VEGFR inhibitor) and their combination.

## Discussion

In this study, we developed an agent-based model to simulate the anti-angiogenesis effect by using VEGFRI treatment in brain tumors. We designed some rules to simulate the tip endothelial cells’ migration, sprout branching and apoptosis based on VEGFR signaling pathways. Together with EGFR signaling pathway equipped in tumor cells considered in our previous studies, we have developed a multiscale agent-based model for angiogenesis-tumor system. Using our model, we could investigate how tumor cells and angiogenesis respond to EGFRI treatment and VEGFRI treatment in a more close-to-reality environment. We revealed a novel angiogenesis-induced drug resistance mechanism, and predicted a synergistic drug combination using EGFR inhibitor and VEGFR inhibitor targeting both tumor cells and angiogenesis, which was consistent with the experimental data.

We further determined the optimal combination timing of EGFRI and VEGRI. The timing of combing VEGFRI was determined to be optimal around 240 hours, which is slightly earlier than the rebound point in the survival rate curve. We could anticipate that adding VEGFRI after the emergence of drug resistance (e.g., after 250 hours) might be too late to rescue the recovering of tumor cells’ survival rate. On the other hand, one may ask whether is it always true that more earlier VEGFRI was added, more benefit we would get? Our simulation demonstrated that adding VEGFRI before 240 hours did have an obvious influence on change of the amount of tumor cells and ECs, however, the tumor cell survival rates were almost surprisingly the same. In addition, compared with the EGFRI-only treatment, we would observe the tumor cells’ survival rates under different schedules coincided during the early phase as well (0-240 hours).

We interpret the above observations as follows. The apoptosis of ECs due to VEGFRI treatment largely affected the supplement of nutrients to tumor cells, resulting in apoptosis and quiescence of tumor cells. However, the lack of nutrients promoted tumor cells release more tumor-induced angiogenesis factor (TAF), such as VEGF, which largely activated the effective amount of VEGFR, enabling VEGFR to bind with TAF and meanwhile decreasing the effect of VEGFRI in the early stage. And then, the VEGFRI treatment had been gradually becoming predominant. With the indeed loss of active tip ECs, the concentration of nutrients was reduced and most of the sprouts stopped branching or migration, which consistently forced the tumor cells to die.

Our model has some limitations which requires further development in our future work. At this phase, we merely considered the timing of coming VEGFRI treatment with EGFRI treatment. In the future work, we will investigate the dose-dependent effect of the drug combination. In addition, there are a lot of factors should be considered under real circumstances, such as permeability of vessel [26] influenced by drug during EGFRI and VEGFRI treatment and tumor interstitial pressure [27].

Furthermore, some drugs, such as vandetanib [28], have been designed to inhibit tumor growth by targeting EGFR and VEGFR at the same time, which have been under the clinical trials. Vandetanib acts as a kinase inhibitor of a number of cell receptors, mainly the VEGFR, EGFR and the RET-tyrosine kinase [29]. This may resist the influence of potential rejection or interaction of different TKI and VEGFRI treatment drugs, which might offer us a new way of suppressing tumor growth and angiogenesis. In our future work, we will investigate the effect of multi-targets drugs based on the signaling network modeling and simulation.

In summary, we developed a single cell-based multi-scale spatio-temporal model to investigate the role of angiogenesis in drug response of brain tumors. The model integrates four scales: molecular scale (EGFR signaling pathway, cell cycle pathway, VEGFR signaling pathway), cellular scale (tumor cells’ phenotype switch and endothelial cells’ migration), micro-environmental scale (growth factors and nutrients) and tissue scale (angiogenesis). The developed multiscale model demonstrated dual roles of angiogenesis in drug treatment of brain tumors and revealed a novel mechanism of angiogenesis-induced drug resistance. Furthermore, the model predicted a synergistic drug combination targeting both EGFR and VEGFR pathways with optimal combination timing. This study has been dedicated to elucidate mechanistic and functional mechanisms of angiogenesis underlying tumor growth and drug resistance, advancing our understanding of drug resistance, and provides implications for designing more effective drug combination cancer therapies.

## Materials and Methods

### Model assumptions

The major assumptions of our model include:

a. Angiogenesis secretes TGFα, glucose and oxygen into the microenvironment, which mediates the EGFR signaling pathways and cell cycle pathways within tumor cells and influences tumor cells’ activities
b. The tumor cells in turn secrete TAF (e.g., VEGF) into the microenvironment, which stimulates the VEGFR signaling pathways, determining the survival or migration of endothelial cells.

### Multiscale modeling of vascular tumor growth

Based on the above biological mechanisms, we developed a single cell-based multiscale spatio-temporal model to simulate vascular tumor growth. Our model encapsulates four biological scales: molecular, cellular, microenvironmental and tissue scales. At molecular scale, EGFR signaling pathway, cell cycle and VEGFR signaling pathway were considered; at cellular scale, tumor cells switch their phenotypes and endothelial cells migrate, proliferate or dead; at microenvironment scale, growth factors, nutrients (glucose and oxygen) and chemical inhibitors diffuse and exchange; at tissue scale, new blood vessels grow and branch to form a micro-vasculature network. Intracellular signaling pathways were described using ODEs, and microenvironmental factors were described with PDEs. The cell’s phenotype switch was simulated using rule-based algorithm that is determined by both signaling pathways and microenvironmental factors. The migration and branching of micro-vasculatures at tissue scale were determined by VEGF chemotaxis and fibronectin/haptotaxis in microenvironment. The treatment effects of EGFR inhibitor and VEGFR inhibitor were integrated into the model based on their action mechanisms of corresponding signaling pathways. The details of the multiscale modeling of vascular tumor growth was provided in **Text S1**.

### Integrating targeted drug treatment

We integrated the targeted drug treatment with EGFR inhibitor (e.g., gefitnib) and VEGFR inhibitor (sorafenib) into the model.

The EGFRI molecules first permeate the blood vessels and then diffuse in the microenvironment, binding to EGFR to inhibit tumor growth accumulatively. We assumed that the number of EGFRI molecules are large enough, which, therefore, can be treated as continuous variable. The concentration of EGFRI ([*I*_1_](*t*, *x*)) can be described using the following partial differential equations (PDEs):

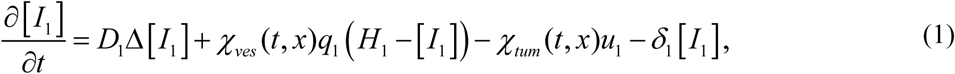

where *D*_1_, *q*_1_, *H*_1,_ *u*_1,_ and *δ*_1_ represent the diffusivity, vessel permeability, in-blood concentration, tumor cell’s uptake rate and natural decay rate of EGFRI, respectively.

Similarly, VEGFRI molecules also permeate through vessels, diffuse in microenvironment, and then bind to the VEGFR and generate accumulative inhibitive effect on vascular growth. The concentration of VEGFRI ([*I*_2_](*t*, *x*)) could be described as follows:

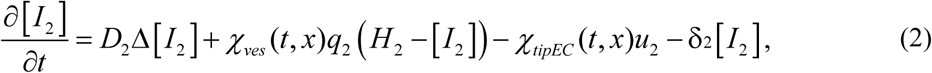

where *D*_2_, *q*_2_, *H*_2_, *u*_2_, and *δ*_2_ represent the diffusivity, vessel permeability, the in-blood concentration, uptake rate by tip endothelial cells and natural decay rate of VEGFRI.

The binding and unbinding processes of inhibitors and receptors are shown below,

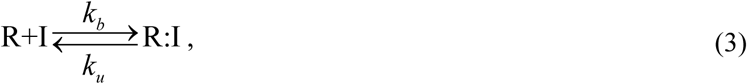

where R represents EGFR or VEGFR, and I represents EGFRI or VEGFRI. We used the Hill function to estimate the concentration of EGFR:EGFRI or VEGFR:VEGFRI complex as follows:

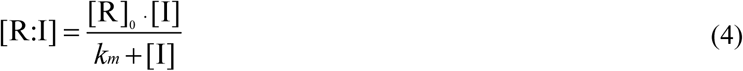

where 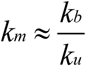 is the Michaelis constant, and [R]_0_ is the initial concentration of EGFR or VEGFR. So, we could derive the amount of effective EGFR and effective VEGFR as follows:

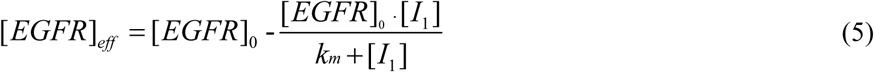

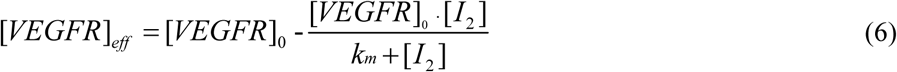

In the above equations, [*I*_1_] and [*I*_2_] at different locations were calculated using Equation (1-2). Therefore, the concentrations of [*EGFR*]_*eff*_and [*VEGFR*]_*eff*_ are spatially heterogeneous.

Under the EGFRI treatment, the effective amount of EGFR of some tumor cells will decrease, resulting in a slow rate of change of the concentration of PLCγ, which largely reduces the migration potential of these tumor cells. See details in **Text S1**.

During the VEGFRI treatment, the amount of effective VEGFR may decrease, which might largely reduce the survival rate of tip endothelial cell, and also the growth, migration or branching. Hence, we set some new rules to simulate endothelial cell’s fate determination. For each tip EC, we first check whether the concentration of effective VEGFR at the current location is higher than the average concentration of VEGFR at the locations of all active ECs. If so, we turn to the sprout migration or branching rules. Otherwise, the tip EC turns to irreversible apoptosis state that cannot migrate or branch any longer. See details in **Text S1**.

### Summary of simulation algorithm

The algorithm iteratively repeated the following steps until the end of the simulation (450 hours):

1. Microenvironmental scale: solve PDEs to calculate the distribution of glucose, O_2_, TGFα, TAF and EGFRI as well as VEGFRI.
2. Molecular scale: solve ODEs to simulate EGFR and cell cycle signaling pathways; integrating EGFRI or VEGFRI to determine effective EGFR and VEGFR.
3. Cellular scale: simulate phenotype switch of tumor cells and endothelial cells.
4. Tissue scale: simulate tip endothelial cells’ apoptosis, migration and sprout branching according to the distributions of VEGF and fibronectin.

### Bliss combination index

The Bliss combination index [30, 31] was calculated as follows:

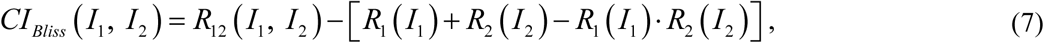

where *I*_1_ and *I*_2_ represent EGFRI and VEGFRI, respectively. *R*_12_ (*I*_1_, *I*_2_) is killing rate of tumor cells by combined inhibitors. *R*_1_ (*I*_1_) and *R*_2_ (*I*_2_) are the killing rates of tumor cells by EGFRI alone or VEGFRI alone, respectively. If the combination index (CI) is greater than 0 then the drug combination has synergistic effect, whereas if CI is less than 0 then the combination effect is antagonistic.

### Survival analysis

The clinical survival data and RNA-seq data of glioma patients were downloaded from TCGA database (https://cancergenome.nih.gov/). The dataset included a total of 648 GBM cases with clinical follow-up information, and 173 GBM cases with level 3 RNA-seq Gene expression data. By matching the sample ID, we got 167 GBM cases with full data of both clinical and gene expression. The gene symbols/aliases coding the proteins in the VEGFR signaling pathways were listed in **Table S6**. COX proportional hazards (PH) regression model [32] was used to compute the risk score predicted by expression of genes related to VEGFR signaling pathways in angiogenesis. The patients were divided into 2 groups according to the median of the risk score. K-M curves were plotted for these two groups of patients, respectively. Log-rank test was used to assess the significance of difference between two survival curves.

## Acknowledgements

X Sun was supported by grants from the National Natural Science Foundation of China (61503419), the Guangdong Nature Science Foundation (2016A030313234), the fund for Guangdong Provincial Key Laboratory of Orthopedics and Traumatology (2016B030301002) and the Opening Project of Guangdong Province Key Laboratory of Computational Science at the Sun Yat-sen University (2018003).

## Authors’ contributions

WL performed simulation, analyzed data and drafted the manuscript. XS designed the study, analyzed data and wrote the manuscript. JZ participated in discussion and data interpretation. All authors read and approved the final manuscript.

## Supporting Information

**Text S1**. Details of multiscale modeling.

**Figure S1**. 3-D vascular tumor profile at 150 hours from different views.

**Figure S2**. The survival rate of tumor cells under treatment of EGFRI combining VEGFRI at different time points before 240 hours.

**Figure S3**. Spatial distributions of concentrations of EGFRI and VEGFRI at the end of the simulation.

**Figure S4**. Spatial distributions of different microenvironment factors under the combined treatment of VEGFRI and EGFRI at the end of the simulation.

**Table S1**. Kinetic equations of EGFR signaling pathway.

**Table S2**. Coefficients of kinetic equations of the EGFR signaling pathway.

**Table S3**. Kinetic equations of the cell-cycle.

**Table S4**. Parameter in cell-cycle pathway.

**Table S5**. Parameters of PDEs in the model.

**Table S6** Genes in VEGFR signaling pathways.

